# Single-sequence protein structure prediction using supervised transformer protein language models

**DOI:** 10.1101/2022.01.15.476476

**Authors:** Wenkai Wang, Zhenling Peng, Jianyi Yang

## Abstract

It remains challenging for single-sequence protein structure prediction with AlphaFold2 and other deep learning methods. In this work, we introduce trRosettaX-Single, a novel algorithm for single-sequence protein structure prediction. It is built on sequence embedding from s-ESM-1b, a supervised transformer protein language model optimized from the pre-trained model ESM-1b. The sequence embedding is fed into a multi-scale network with knowledge distillation to predict inter-residue 2D geometry, including distance and orientations. The predicted 2D geometry is then used to reconstruct 3D structure models based on energy minimization. Benchmark tests show that trRosettaX-Single outperforms AlphaFold2 and RoseTTAFold on natural proteins. For instance, with single-sequence input, trRosettaX-Single generates structure models with an average TM-score ~0.5 on 77 CASP14 domains, significantly higher than AlphaFold2 (0.35) and RoseTTAFold (0.34). Further test on 101 human-designed proteins indicates that trRosettaX-Single works very well, with accuracy (average TM-score 0.77) approaching AlphaFold2 and higher than RoseTTAFold, but using much less computing resource. On 2000 designed proteins from network hallucination, trRosettaX-Single generates structure models highly consistent to the hallucinated ones. These data suggest that trRosettaX-Single may find immediate applications in de novo protein design and related studies. trRosettaX-Single is available through the trRosetta server at: http://yanglab.nankai.edu.cn/trRosetta/.

## Introduction

AlphaFold2 ^1^ and other protein structure prediction methods, such as RoseTTAFold ^2^, trRosetta ^3^ and trRosettaX ^4^, make use of the co-evolution signal embedded in a pre-generated multiple sequence alignment (MSA). However, no MSA could be built for proteins that do not have any homologous sequences in the current sequence database. In our test with the CASP14 targets, all methods perform poorly for single-sequence input (see Figure S1). For example, the average TM-score ^5^ for the models predicted by AlphaFold2 drops dramatically from >0.8 (with MSA) to ~0.3 (with single sequence). Interestingly, all tested methods (AlphaFold2, RoseTTAFold and trRosettaX) show similar level of accuracy for single-sequence input. We conclude that it remains challenging to predict accurate structure with single-sequence information, even with the state-of-the-art methods. In practice, there do exist some proteins (e.g., from eukaryote) with limited number of homologous sequences. It is thus worthwhile developing single-sequence protein structure prediction methods.

Many protein language models ^6–11^ have been developed in recent years, inspired by the development of new natural language processing approach, especially transformer ^12^ and BERT ^13^. These models are typically trained on large sequence database in an unsupervised way, i.e., to generate training objectives from the sequences alone. Subsequent small-scale supervised training for downstream tasks, e.g., the prediction of secondary structure and inter-residue contact, shows that the pre-trained models are helpful for these structure-related tasks even if they are trained with sequence information only ^6^. These successes pave the way for developing accurate deep learning-based approach to single-sequence protein structure prediction.

Compared with MSA-based protein structure prediction, only limited studies were done for single-sequence protein structure prediction with deep learning. SSCpred ^14^ is a deep convolutional network for contact map prediction using sequence one-hot encoding and 23 predicted 1D structural features. It is improved by SPOT-Contact-Single ^15^ by using a pre-trained language model ESM-1b ^6^. Both SSCpred and SPOT-Contact-Single predict the 2D contact map only. To the best of our knowledge, RGN2 is the first reported deep learning-based single-sequence method for 3D structure prediction ^16^. RGN2 makes use of a transformer protein language model to learn structural information and uses a geometric module to generate the backbone structure. However, no web server or standalone package is available for RGN2.

In this work, we introduce trRosettaX-Single, a deep learning-based single-sequence protein structure prediction method with supervised transformer protein language model. Benchmark tests show that our method outperforms AlphaFold2 and RoseTTAFold on natural proteins. On designed proteins, trRosettaX-Single is competitive with AlphaFold2 and outperforms RoseTTAFold. trRosettaX-Single also generates significantly more accurate contact prediction than SPOT-Contact-Single on all independent test sets.

## Results

### Overview of trRosettaX-Single

The overall architecture of trRosettaX-Single is shown in Figure 1A. The only input to trRosettaX-Single is the amino acid sequence of a target protein. The sequence is fed into a new transformer protein language model s-ESM-1b to obtain single representation and attention maps (pair representations). Together with one-hot encoding, the protein sequence is represented as a *L*×*L*×4756 tensor, which is the input to a multi-scale network (denoted by Res2Net_Single) used in trRosettaX. The network outputs the predicted 2D geometry, including inter-residue distance and orientations defined in trRosetta^3^. The predicted 2D geometry is then converted into spatial constraints to guide structure folding based on fast energy minimization.

**Figure 1.**
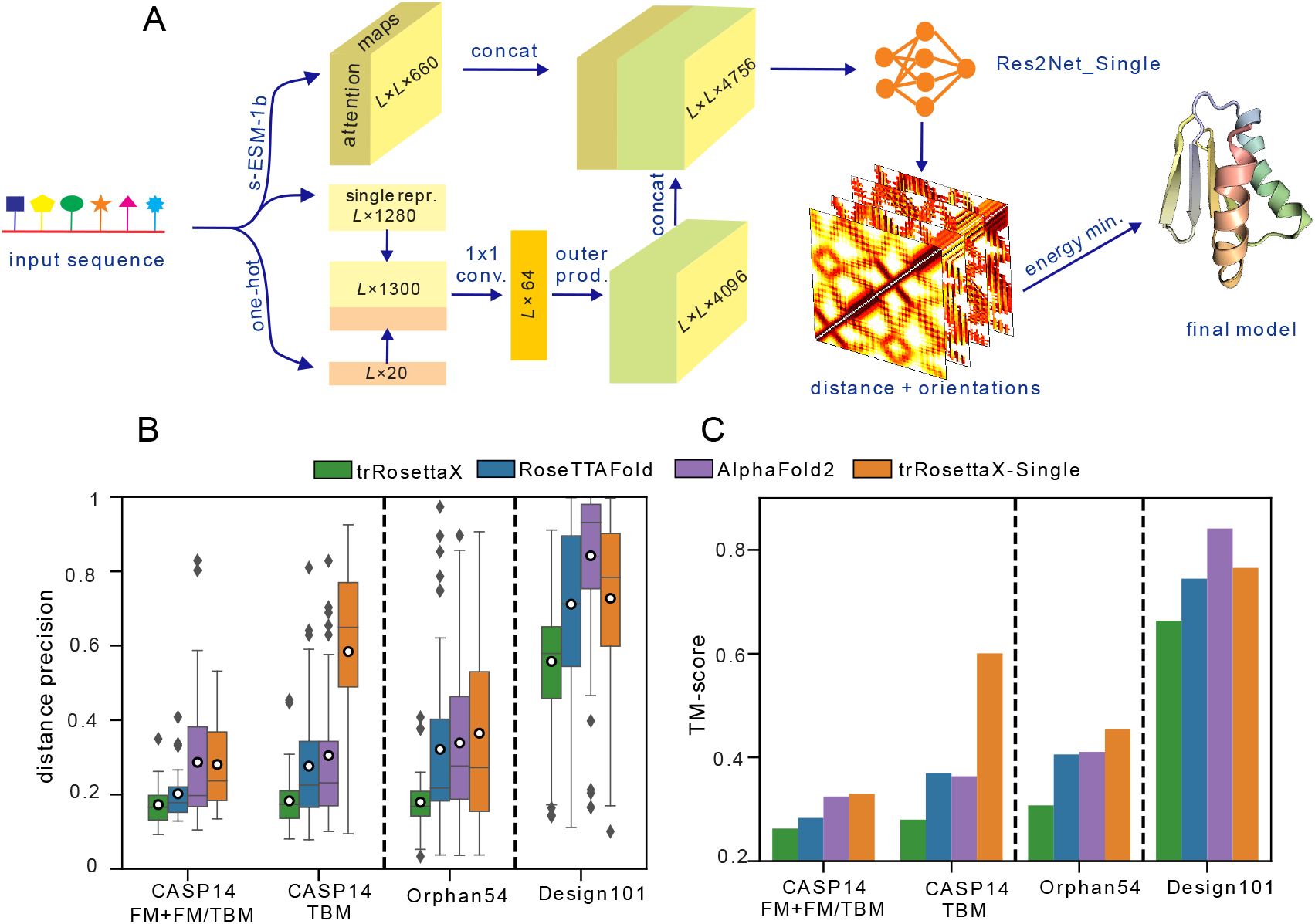
The architecture and performance of trRosettaX-Single. A. overview of trRosettaX-Single. s-ESM-1b is a supervised transformer protein language model with initial parameter from ESM-1b. Res2Net_Single is a knowledge-distilled multi-scale neural network. B. comparison with MSA-trained methods in terms of the precision of predicted inter-residue distances. The bottom and top of each box refer to the first and third quartiles, respectively. The horizontal line and the white hole inside the box refer to the median and the mean, respectively. C. comparison with MSA-trained methods in terms of the average TM-score of the predicted structure models. All predictions are made without any sequence or structural homologs.

### Comparison with MSA-trained methods

We compare trRosettaX-Single with three methods trained with MSA (AlphaFold2, RoseTTAFold and trRosettaX) in terms of accuracy of predicted inter-residue distance and structure models on three benchmark datasets. The accuracy of predicted inter-residue distance is measured by *distance precision* proposed in ^17^. The residue pairs are first ranked by the predicted probability of inter-residue distance ≤ 20 Å. The distance precision is then defined as the ratio of *correctly predicted* residue pairs (i.e., the difference between the predicted and the real distances is less than 2 Å) over the top 15*L* residue pairs (separation ≥ 12), where *L* is the length of sequence. The accuracy of the predicted structure models is measured by TM-score ^5^. All methods are installed and run locally without any sequence or structural homologs. For RoseTTAFold, we only assess its pyRosetta version, which was more accurate than its e2e version in our observations.

#### Performance on the CASP14 dataset

The full-length sequences for 51 targets from CASP14 are first submitted to all methods to predict inter-residue distances/contacts and full-length structure models. Similar to the official evaluation, the predictions are then trimmed into domains based on the official domain definitions for the subsequent domain-based assessment. On CASP14’s 77 domains, trRosettaX-Single achieves a mean distance precision of 0.475 for the predicted inter-residue distances, significantly higher than AlphaFold2 (0.298), RoseTTAFold (0.249) and trRosettaX (0.179) (Table S1). Figure 1B shows that all methods have higher distance precision for the TBM domains than the FM+FM/TBM domains, though no sequence or structure homologs were used for both types of domains. This may suggest that the TBM domains are on average easier to fold than the FM+FM/TBM domains by nature, as further illustrated by the quality of the predicted structure models. trRosettaX-Single’s average TM-score is 0.596 for 50 TBM domains (Figure 1C), for which 33 have correctly predicted fold (i.e., TM-score > 0.5). In comparison, the average TM-scores (resp. the number of domains with correctly predicted fold) are 0.364, 0.371, 0.28 (resp. 10, 10, 4) for AlphaFold2, RoseTTAFold and trRosettaX, respectively (Figure 2A). It remains challenging to predict the structure for CASP14’s FM+FM/TBM domains with single sequence, for which trRosettaX-Single and AlphaFold2 have similar average TM-score (~0.33), much lower than the TBM domains.

**Figure 2.**
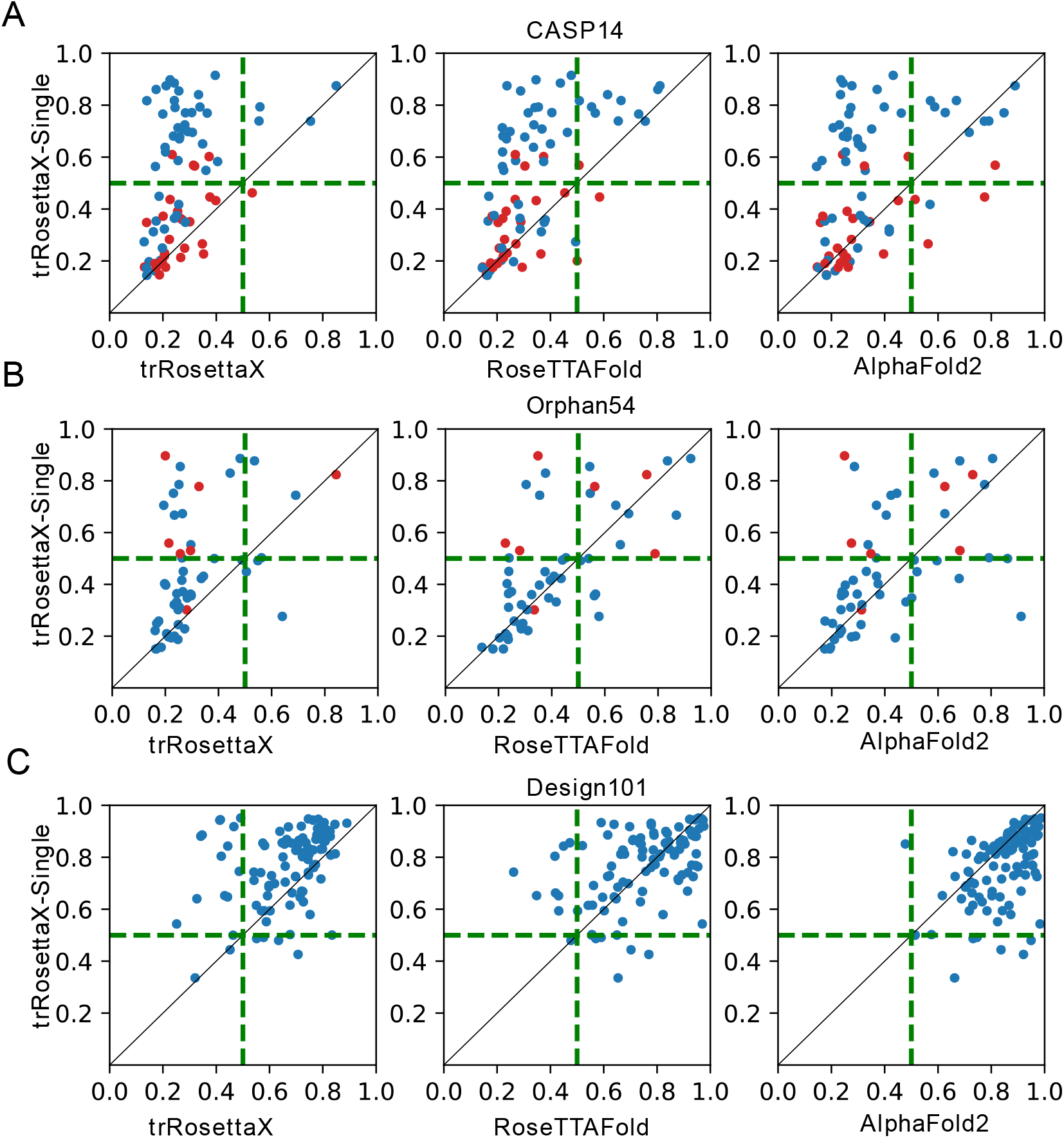
Head-to-head TM-score comparison between trRosettaX-Single and other MSA-trained methods. The red and blue points in A are for CASP14 FM+FM/TBM and TBM domains, respectively. The red points in B are targets without any sequence homologs. The dashed horizontal and vertical lines in A-C correspond to TM-scores of 0.5.

#### Performance on targets with few sequence homologs

As there exists a significant number of sequence homologs for most CASP14 targets (even for the FM targets), we further tested all methods on a set of 54 targets (named as Orphan54) with limited number of homologous sequences in the current sequence database (the effective number of homologous sequences is less than 10 according to HHblits ^18^ search against the Uniclust30_2018). Figure 1B shows that the average precision of the predicted distances on this dataset by trRosettaX-Single (0.365) is higher than AlphaFold2 (0.339), RoseTTAFold (0.321) and trRosettaX (0.179). When this improved distance is used in the subsequent folding, trRosettaX-Single generates more accurate structure models than others: the average TM-scores (Figure 1C) are 0.455, 0.411, 0.406, 0.307 for trRosettaX-Single, AlphaFold2, RoseTTAFold and trRosettaX, respectively.

A total of 7 proteins in Orphan54 do not have any homologous sequences in Uniclust30_2018 (red points in Figure 2B). For these proteins, the average TM-score of the structure models predicted by trRosettaX-Single is 0.63, significantly higher than AlphaFold2 (0.46), RoseTTAFold (0.471) and trRosettaX (0.345). trRosettaX-Single is able to predict correct fold for six out of these proteins. Figure 3A shows the results on a representative target (PDB ID: 6LF2). The distance maps predicted by AlphaFold2 and RoseTTAFold are blurred with low distance precisions (0.178 and 0.326, respectively). The predicted structure models also have low TM-scores, i.e., 0.248 for AlphaFold2 and 0.348 for RoseTTAFold. In contrast, the distance map predicted by trRosettaX-Single is similar to the native distance map with a distance precision of 0.906; and the predicted structure model has a high TM-score of 0.897.

**Figure 3.**
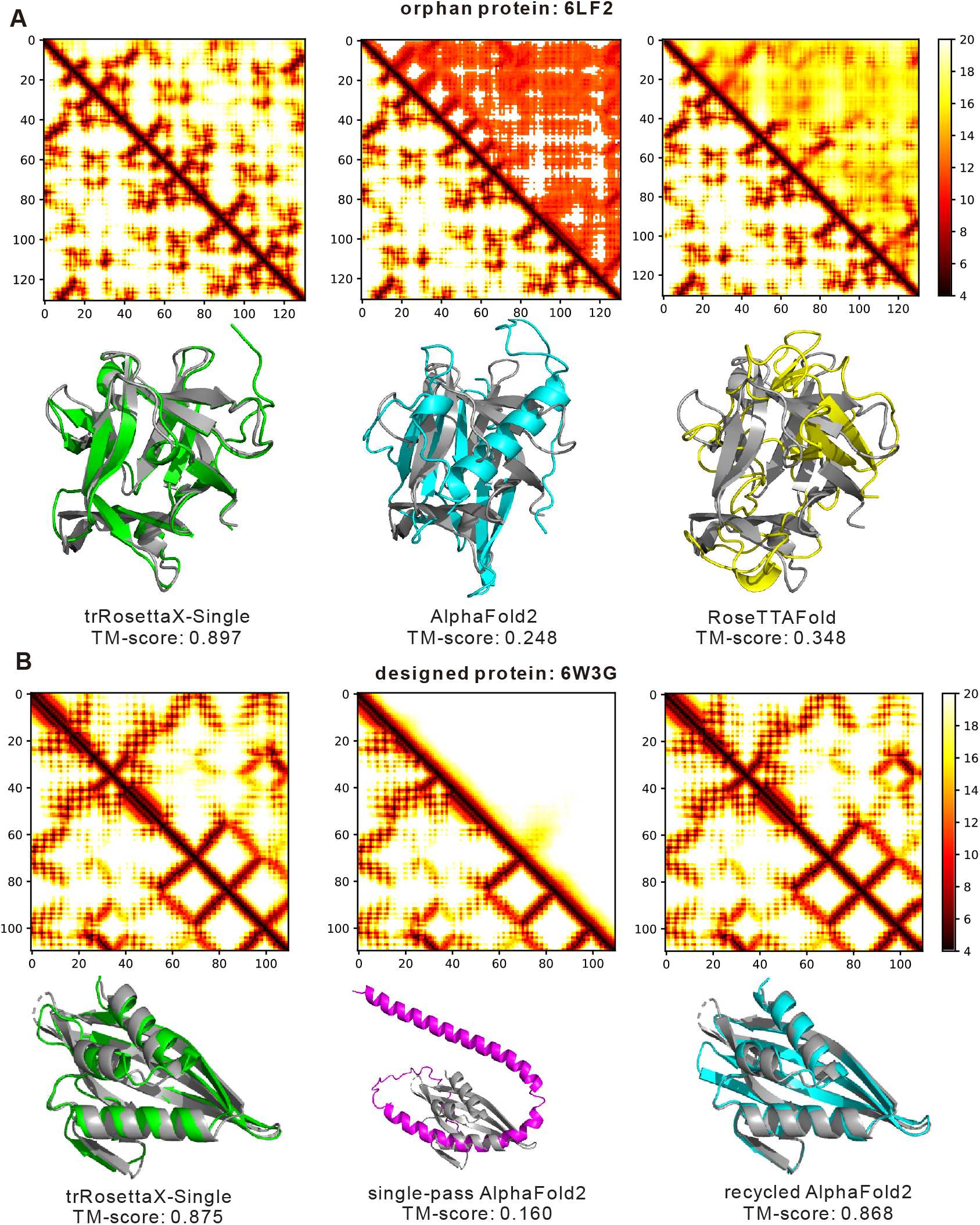
Comparison between trRosettaX-Single, AlphaFold2 and RoseTTAFold on two example proteins. The lower and upper triangles are the native and predicted distance maps, respectively. The predicted structure models and the native structure are shown in color and gray cartoons, respectively. A. comparison with AlphaFold2 and RoseTTAFold on 6LF2, an orphan protein without any homologous sequence. B. comparison with both the single-pass and the recycled AlphaFold2 on a human-designed protein 6W3G.

#### Performance on human-designed proteins

Human-designed proteins are ideal candidates for benchmarking single-sequence folding. Here we evaluate our method on 101 human-designed proteins. Figures 1B, 1C and Figure 2 show that all methods predict much more accurate inter-residue distances and structure models on these proteins. For example, trRosettaX-Single achieves a mean TM-score of 0.765 on this dataset, significantly higher than that on other datasets (0.455 on Orphan54; 0.497 on CASP14 domains). This is consistent with our previous observation that trRosetta generates significantly more accurate structures for designed proteins than natural proteins^3^. This is probably because the designed proteins have been manually optimized with exceptional stability and easier to predict than natural proteins. Except AlphaFold2, trRosettaX-Single outperforms RoseTTAFold and trRosettaX on the designed proteins. Figures 1C and 2C show that trRosettaX-Single achieves an average TM-score of 0.765 and generates correct fold for 95 out of 101 designed proteins, higher than RoseTTAFold (0.745, 92) and trRosettaX (0.663, 85).

We find that the recycle mechanism in AlphaFold2 plays a key role in improving the structure models for designed proteins. If we run AlphaFold2 with no recycle (i.e., walking through the whole model only once, by assigning the parameter “num_cycle” to zero), the average TM-score drops from 0.841 to 0.649, lower than trRosettaX-Single (Figure S2). Figure 3B shows a representative example (PDB ID: 6W3G), on which the TM-scores of the models predicted by the single-pass and the recycled AlphaFold2 are 0.16 and 0.868, respectively. The predicted structure model by the single-pass AlphaFold2 is a turned alpha-helix with no long-range interactions. The recycle mechanism helps AlphaFold2 to fix the interactions, resulting in a much more accurate predicted structure. trRosettaX-Single achieves a TM-score of 0.875 on this protein, which is significantly higher than the single-pass AlphaFold2 model and close to the recycled one. Inspired by AlphaFold2, we also tried to train a model with similar mechanism but did not see significant improvement. This may be because our method is not end-to-end and only the predicted 1D and 2D information rather than 3D structure information could be fed into the network.

### Comparison with methods trained with single sequence

As mentioned in the Introduction, a few methods trained with single sequence have been reported in the literature, including SSCpred, RGN2 and SPOT-Contact-Single ^15^. As RGN2 is not available and SPOT-Contact-Single was shown to outperform SSCpred, we compare trRosettaX-Single with SPOT-Contact-Single only. The comparison is based on contact precision because SPOT-Contact-Single predicts inter-residue contacts rather than 3D structure. The contact precision is defined as the number of correctly predicted contacts out of the predicted top *L* long-range contacts (i.e., with distance ≤ 8 Å and sequence separation ≥ 24). The mean precisions of the predicted contacts by both methods are shown in Figure 4A, suggesting that trRosettaX-Single consistently outperforms SPOT-Contact-Single for all benchmark datasets. Head-to head comparisons in Figure 4B indicate that for most targets, trRosettaX-Single achieves more accurate contact predictions than SPOT-Contact-Single. For example, the average precisions of trRosettaX-Single and SPOT-Contact-Single on the 101 human-designed proteins are 0.608 and 0.41, respectively. For 92 out of these proteins, trRosettaX-Single has higher precision than SPOT-Contact-Single. As both methods use protein language models to encode the single sequence, the superior performance of trRosettaX-Single over SPOT-Contact-Single may be attributed to the more powerful multi-scale network Res2Net (compared with ResNet in SPOT-Contact-Single) and a few key factors, such as the supervised training of the pre-trained ESM-1b, knowledge distillation, etc. More detailed discussions about these factors are given below.

**Figure 4.**
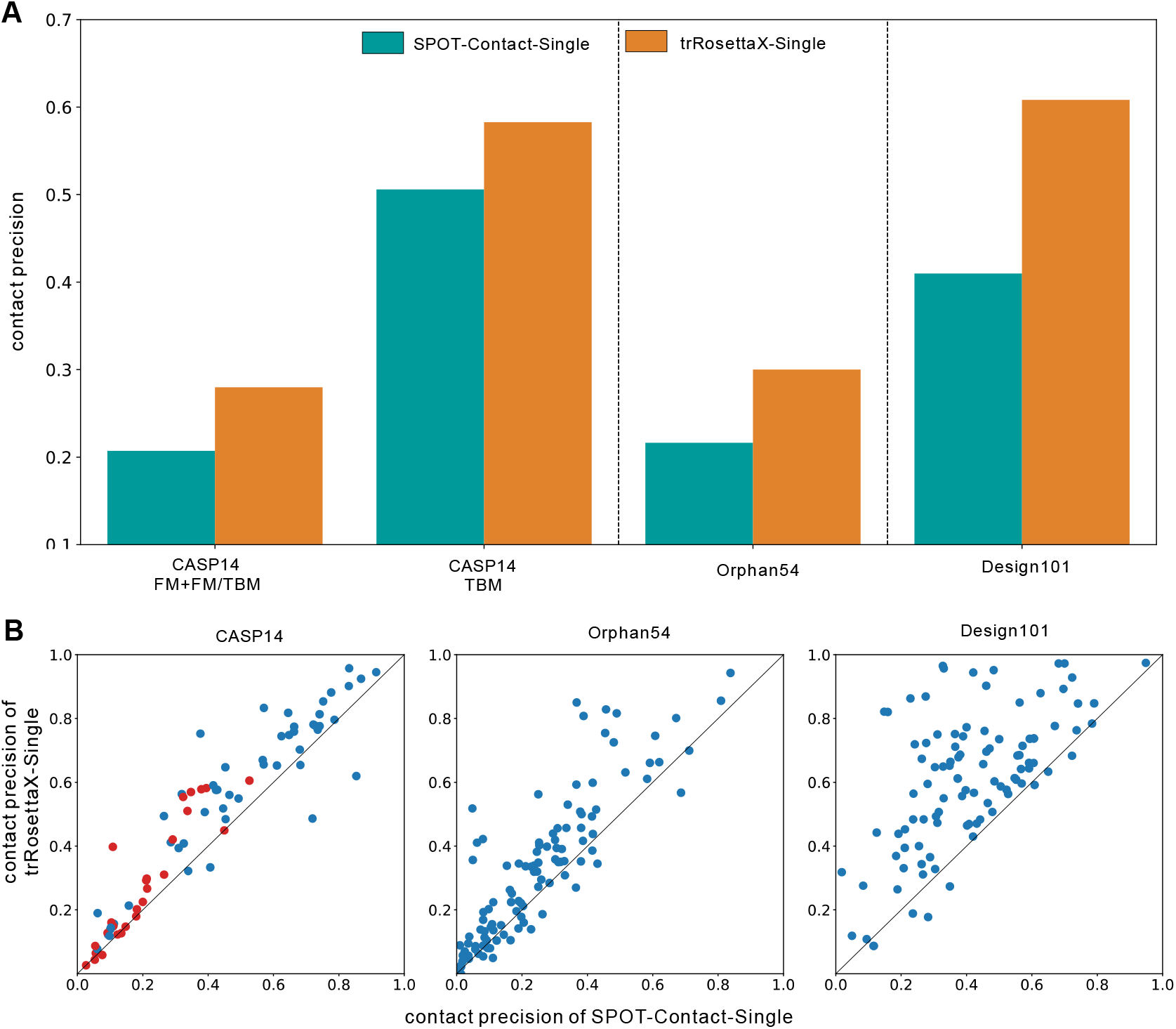
Precision of predicted contacts. A. average precisions of the top *L* long-range contacts predicted by trRosettaX-Single and SPOT-Contact-Single on the benchmark datasets. B. head-to-head comparisons based on contact precision. The red and blue points in the first plot are CASP14 FM+FM/TBM and TBM domains, respectively.

### Application to hallucinated proteins

We further test our method on the 2000 hallucinated proteins, which were *de novo-designed* by deep network hallucination^19^. As shown in Figure 5A, the predicted structure models by trRosettaX-Single are highly consistent to the hallucinated structures, with a mean TM-score of 0.902. As the experimental structures for most of these proteins are unknown, we estimate the TM-score of the predicted models (see the next section). The average of the estimated TM-scores of the predicted structure models for these proteins is 0.86. For all proteins, the predicted models are estimated to have the correct fold. On three proteins (0217, 0515 and 0738) that have been determined by X-ray diffraction or NMR experiments, trRosettaX-Single generates structure models with similar accuracy (i.e., TM-score and RMSD) to the hallucinated ones (Figure 5B). These data illustrate again the potential application of trRosettaX-Single in protein design. The high accuracy achieved on designed/hallucinated proteins implies the possibility of developing similar hallucination methods based on trRosettaX-Single.

**Figure 5.**
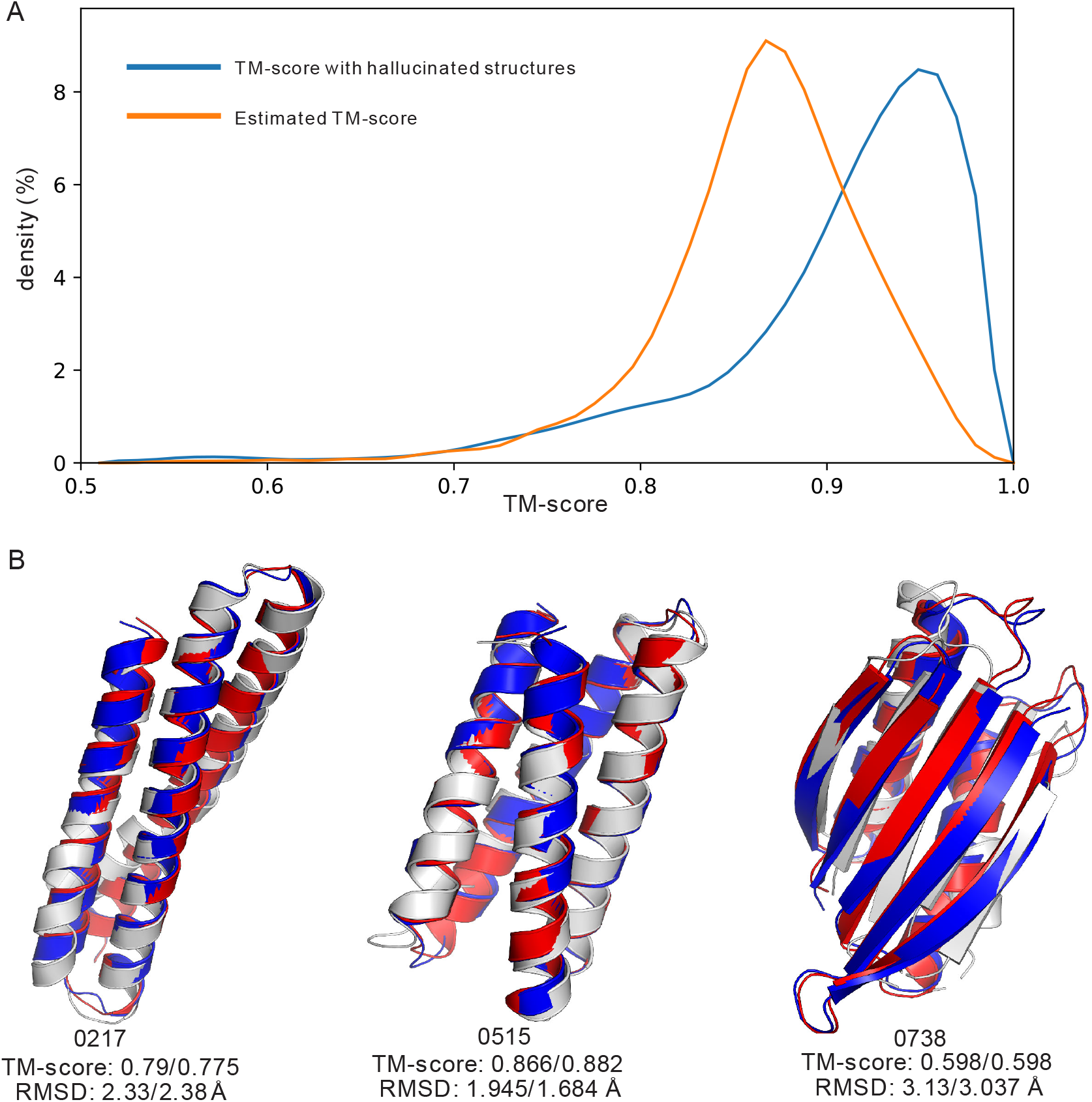
Application to hallucinated proteins. A. TM-score distributions. The blue curve is for the TM-scores between trRosettaX-Single models and the hallucinated models (predicted by trRosetta). The orange curve is for the estimated TM-scores of the trRosettaX-Single models. B. the superposition of the trRosettaX-Single models (blue cartoon) and the hallucinated models (red cartoon) against the experimental structures (grey cartoon) for three hallucinated proteins.

### Ablation study

With supervised learning, we re-trained the language model ESM-1b from its initial parameters. The new model (s-ESM-1b) was then used to generate extra features from single sequence. In addition, a few training strategies were explored to make full use of the limited sequence information (see Methods). To analyze their contributions, we train and evaluate six ablation models below (the datasets used by each model are indicated in parentheses).

a. baseline model using sequence one-hot encoding only (Single15015);
b. baseline + ESM-1b (Single15015);
c. baseline + ESM-1b + knowledge distillation (MSA15015 + Single15015);
d. baseline + s-ESM-1b (Single15015);
e. baseline + ESM-1b + extended training set (Cluster22503);
f. final model with all components listed above (MSA15015 + Single15015 + Cluster22503).

The above ablation models are used to predict the inter-residue distances. trRosettaX is used as a control here as it adopts a similar neural network architecture. To save time, no ensemble is applied and no structure modeling step is performed in this analysis. The differences between the precisions of the predicted distances by the ablation models and trRosettaX are summarized in Figure 6A. When no pretrained language model is used, the baseline model has similar or lower precision than trRosettaX. With the introduction of the ESM-1b features, the predicted distances for natural proteins become much more accurate than trRosettaX. However, for designed proteins, we do not see any significant difference between using or not using ESM-1b. With supervised training in s-ESM-1b, knowledge distillation and extended training set, we are able to make consistent improvements for all datasets. The most accurate model is obtained by considering all components in the final model, which has 0.3, 0.18, 0.21 higher distance precision than trRosettaX on the CASP14, Orphan54 and Design101, respectively.

**Figure 6.**
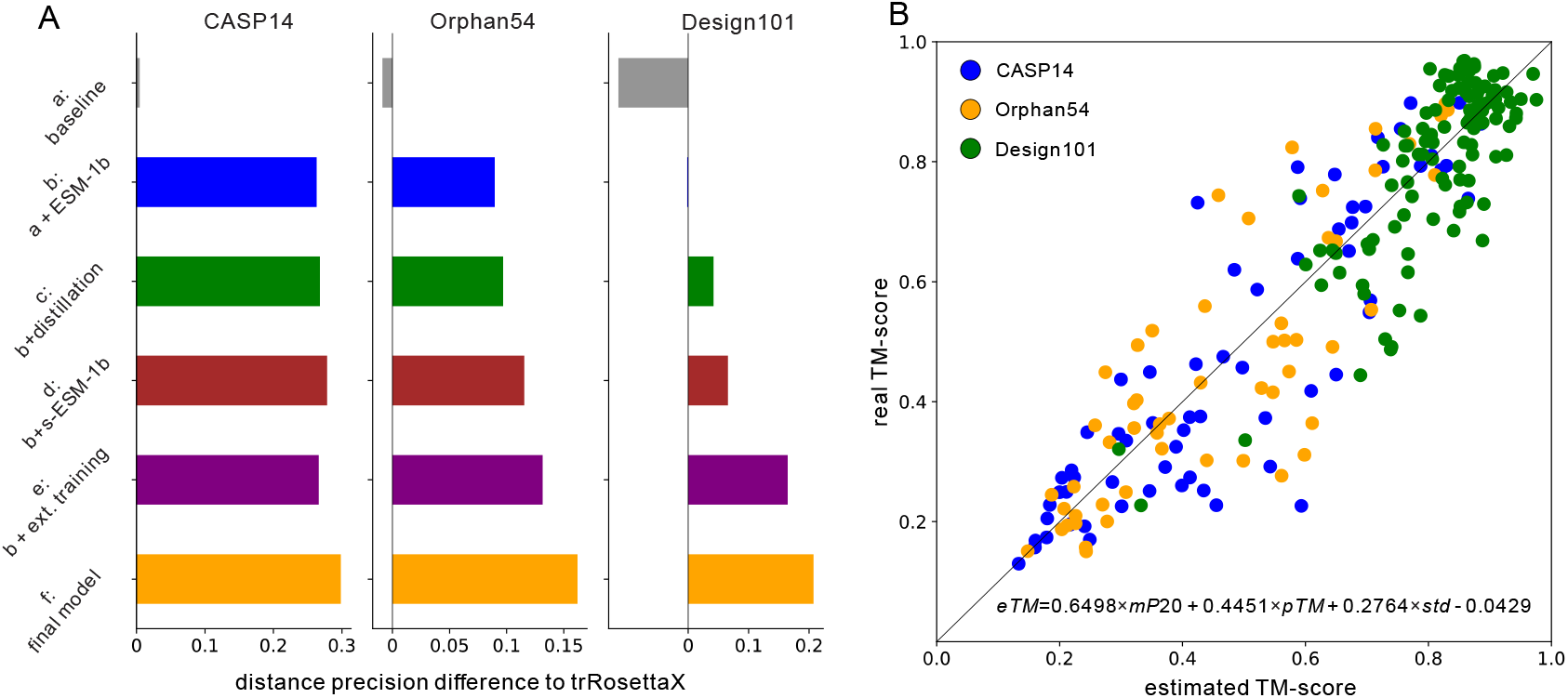
Ablation study and estimation of model accuracy. A. distance precision difference between trRosettaX and other ablation models. B. correlation between the real and the estimated TM-scores. The formula for estimating the TM-score is given at the bottom of the figure.

### Confidence score of predicted structure models

In trRosetta and trRosettaX^3, 20^, the TM-scores of the predicted structure models are estimated using the confidence of the predicted distance and the convergence of the top structure models. Here we extend this estimation in trRosettaX-Single using three variables. The first one is *mP20*^17^, which measures the average probability of the predicted top *15L* inter-residue distances (distance ≤ 20 Å). For each residue pair, the standard deviation of the probability values from the predicted distance distribution is first calculated. The second one is the average of the standard deviations over all residue pairs. The third one is the average pair-wise TM-score of the top 10 non-constrained structure models (the same as in trRosettaX). Linear regression over these three variables is employed to estimate the TM-score of a predicted structure model. For the targets from the three benchmark datasets, the estimated TM-score correlates very well with the real TM-score of the predicted models (Pearson’s *r* is 0.916, Figure 6B).

## Conclusions

We have introduced trRosettaX-Single for single-sequence protein structure prediction using supervised transformer protein language models and distilled multi-scale networks. trRosettaX-Single outperforms AlphaFold2 and RoseTTAFold on natural proteins. trRosettaX-Single achieves an average TM-score of 0.77 on 101 human-designed proteins, which is competitive with AlphaFold2 (0.84) and more accurate than RoseTTAFold (0.76), but with much less computer resource. Compared with the single-sequence-trained contact prediction method SPOT-Contact-Single, trRosettaX-Single is significantly more accurate for all benchmark datasets. The above data show that it is possible to increase the accuracy of single-sequence protein structure prediction with supervised transformer-based protein language models. However, we admit that the accuracy of single-sequence structure prediction for natural proteins is still far from satisfactory. In future we hope to move the single-sequence accuracy towards the MSA-based level further.

## Methods

### Datasets

Two datasets are used to train our network. The first is a high-quality dataset from our previous works^3, 4^, including 15051 non-redundant (< 30% pairwise sequence identity) chains from PDB released before 2018-05. The structures in this dataset are from high-resolution (≤ 2.5 Å) X-ray entries and each chain’s MSA has at least 100 homologous sequences. Knowledge distillation is done with the MSAs in this dataset. For convenience, we denote this dataset by **MSA15015** or **Single15015**, respectively, depending on if MSAs or single sequences are used during training. The second set is an extended version of the first one by relaxing the criteria (i.e., no requirements of structure determination methods and number of homologous sequence). It contains 330080 protein chains released before 2018-05, which are then clustered at 30% sequence identity cutoff, resulting in 22503 clusters. For convenience, this dataset is denoted by **Cluster22503**. At each training epoch, we cycle through all clusters and randomly select a protein chain from each cluster.

Three independent test sets are used to compare our method with others. 1) **CASP14 targets.** We collect 77 domains from CASP14, after removing two targets without experimental structures (T1085 and T1086, involving 6 domains). T1044 (due to its huge size) and its domains (T1031, T1033, T1035, T1037, T1039, T1040, T1041, T1042, T1043) are also removed. 2) **Orphan54**. This set contains 54 non-redundant (< 30% pairwise sequence identity) single sequences from native proteins released after 2020-05 with *N*_eff_ (i.e., the effective number of homologous sequences in MSA) less than 10 (according to HHblits^18^ search against Uniclust30_2018 with default parameters). 3) **Human-designed proteins (Design101).** From PDB, we collect all single-chain structures with keywords “*de novo* designed” or “computational designed” in the structure titles. Structures with <50 or >300 AAs or with too simple topologies (e.g., a single *α*-helix) are removed. Then we run HHblits against Uniclust30_2018 and remove the structures with sequence homologs. The remaining targets are then merged with the 35 designed proteins from previous works^3, 21^, resulting in 101 human-designed targets. Details about the above datasets are summarized in Table S3.

### Network architecture

As shown in Figure S3, trRosettaX-Single’s network (denoted by Res2Net_Single) contains two groups of Res2Net blocks, which output 128 and 256 feature maps, respectively. After the last Res2Net block, four classifiers consisting of a 1×1 convolutional layer and a softmax operation are used to predict the probability distributions of the inter-residue geometries (Cβ-Cβ distance and three orientations, defined in trRosetta ^3^).

### Supervised transformer protein language model s-ESM-1b

The features extracted from unsupervised pre-trained protein language model (i.e., ESM-1b^6^) show strong correlation with some structural characteristics, such as secondary structure, inter-residue contact and ligand-binding site. We propose that the correlation can be further enhanced by supervised training of ESM-1b on specific tasks starting from the pre-trained parameters.

In this work, we re-train the ESM-1b parameters based on supervised learning, resulting in a new model s-ESM-1b (Figure S4). As shown in Figure S4, we optimize ESM-1b on two objectives. The first is to predict the amino acid types of the randomly masked positions (with 15% rate), supervised by the cross-entropy loss (*L_mask_*) between the predicted probability distributions and the one-hot encoding of real types. Note that the calculation of *L_mask_* involves all positions of the sequence to guarantee that the amino acid types of the unmasked positions are also predicted correctly. The second is to predict the inter-residue geometry. The attention maps and the 1D representation of the masked sequence are fed into the network Res2Net_Single together with the one-hot encoding of the predicted sequence to predict the inter-residue geometry, supervised by its cross-entropy loss with the native (*L_geometry_*, the same as those defined in trRosetta). The parameters in Res2Net_Single are also updated in this process. The total loss is *L_mask_* + *L_geometry_* with equal weights.

### Input features

The input to the network includes 1D and 2D features. The 1D features include the one-hot encoding of each residue’s amino acid type (20 channels) and the sequence representation vector (1280 channels) from s-ESM-1b. A linear layer with 1×1 convolution is first used to reduce the number of 1D channels from 1300 (=20+1280) to 64. They are then converted to 4096 (=64^2^) 2D feature maps with outer product operation. Additionally, we extract the attention maps from all 33 layers (20 heads per layer) of s-ESM-1b, resulting in 33×20=660 attention maps. To summarize, the input to Res2Net blocks consists of 4756 (=4096+660) 2D feature maps.

### Knowledge distillation guided by MSA-based network

Knowledge distillation^22^ is a training technique to transfer the knowledge from a confident pretrained network (also named as teacher network) into a pre-mature network (also named as student network). The student network is trained under the supervision of the soft labels generated by the teacher network. In this work, the knowledge from a pre-trained MSA-based network (i.e., the teacher network, denoted by Res2Net_MSA; the MSA features are the same as those used in trRosetta) is distilled to Res2Net_Single (i.e., the student network). During training, the MSA of a training target is fed into the pre-trained teacher network Res2Net_MSA to produce a probability distribution. The Kullback-Leibler divergence between this probability distribution and the one from the student network Res2Net_Single are used as an extra loss (*L*_distill_) together with the geometry loss (*L*_geometry_). Equal weights are used for both losses. The training sets MSA15051 and Single15051 are used as MSAs are needed for the teacher network.

### Training of the final models

The training procedure for building the final model consists of two stages. First, we train a Res2Net_Single model on MSA15051, with the distillation guided by a pre-trained network Res2Net_MSA. The distilled Res2Net_Single parameters are refined by the subsequent training, including the supervised re-training of ESM-1b and the update of Res2Net_Single with extended training set Cluster22503. A total of 6 models are trained with same configurations. The final prediction is based on the ensemble of these models.

