## supporting tables and figures for "Single-sequence protein structure prediction using supervised transformer protein language models"

### Supplementary Material

**Table S1**. **Precision of the predicted inter-residue distances/contacts.** Note that only contact precision is shown for SPOT-Contact-Single because it predicts inter-residue contacts only.

| Method | CASP14 | | | Orphan | *De novo* |
|---|---|---|---|---|---|
| | All (77) | FM+FM/TBM (27) | TBM (50) | (54) | (101) |
| trRosettaX | 0.179/0.194 | 0.170/0.149 | 0.184/0.220 | 0.179/0.145 | 0.558/0.469 |
| RoseTTAFold | 0.249/0.237 | 0.197/0.158 | 0.277/0.282 | 0.321/0.270 | 0.711/0.582 |
| AlphaFold2 | 0.298/0.204 | **0.287**/0.172 | 0.305/0.221 | 0.339/0.240 | **0.842/0.724** |
| SPOT-Contact-Single | 0.401 | 0.207 | 0.505 | 0.221 | 0.41 |
| trRosettaX-Single | **0.475/0.479** | 0.278/**0.280** | **0.58/0.578** | **0.365/0.305** | 0.727/0.608 |

**Table S2. TM-score of the predicted structure models.**

| Method | CASP14 | | | Orphan | *De novo* |
|---|---|---|---|---|---|
| | All (77) | FM+FM/TBM (27) | TBM (50) | (54) | (101) |
| trRosettaX | 0.268 | 0.252 | 0.280 | 0.307 | 0.663 |
| RoseTTAFold | 0.336 | 0.277 | 0.371 | 0.406 | 0.745 |
| AlphaFold2 | 0.347 | 0.325 | 0.364 | 0.411 | **0.841** |
| trRosettaX-Single | **0.497** | **0.325** | **0.596** | **0.455** | 0.765 |

**Table S3. Details of datasets used in this work.**

| Dataset | Usage | Size |
|---|---|---|
| MSA15051 | training | 15051 |
| Single15051 | | 15051 |
| Cluster22503 | | 330080 |
| CASP14 | test | 51 (77 domains) |
| Orphan54 | | 54 |
| Design101 | | 101 |

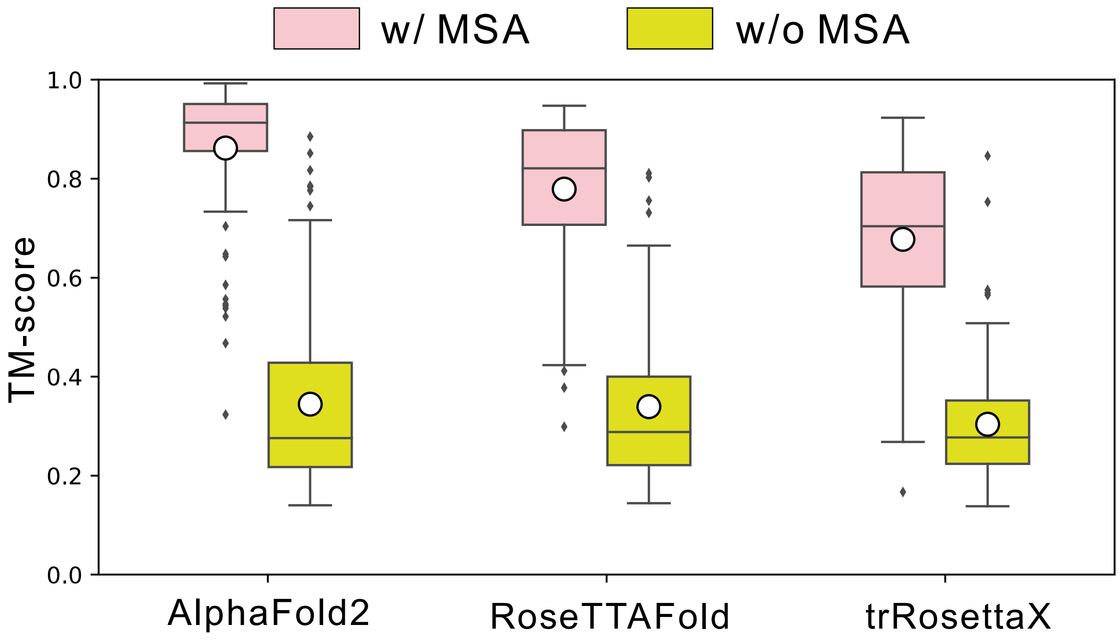


**Figure S1. Comparison between the MSA-based and single-sequence predictions by AlphaFold2, RoseTTAFold and trRosettaX on the CASP14 dataset.** The bottom and top of box refer to the first and third quartiles, respectively. The horizontal line and the white hole inside each box refer to the median and mean, respectively.

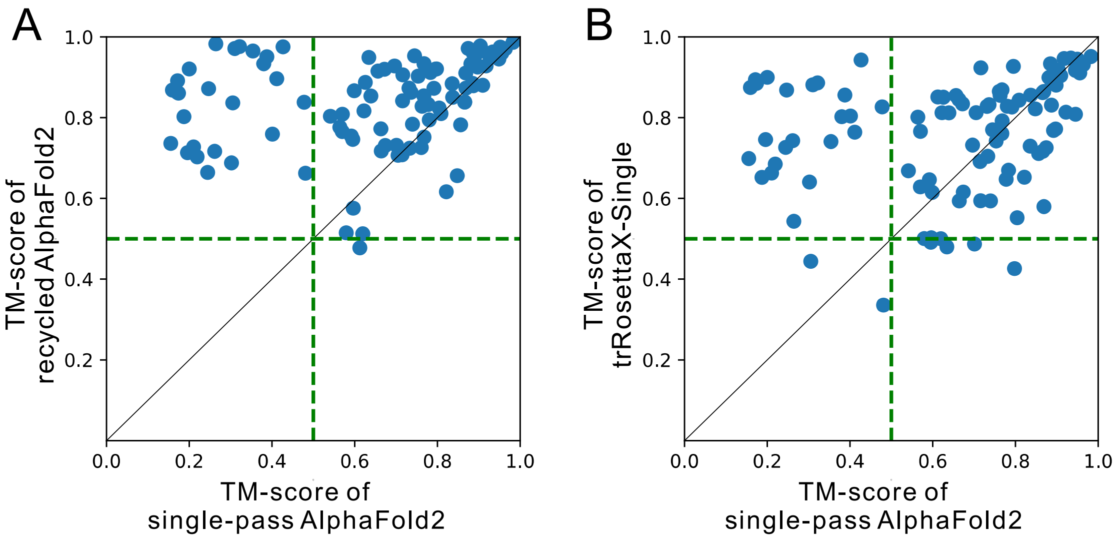


**Figure S2. Impact of recycle to AlphaFold2 on the dataset Design101.** A. the recycled and the single-pass AlphaFold2. B. trRosettaX-Single and the single-pass AlphaFold2.

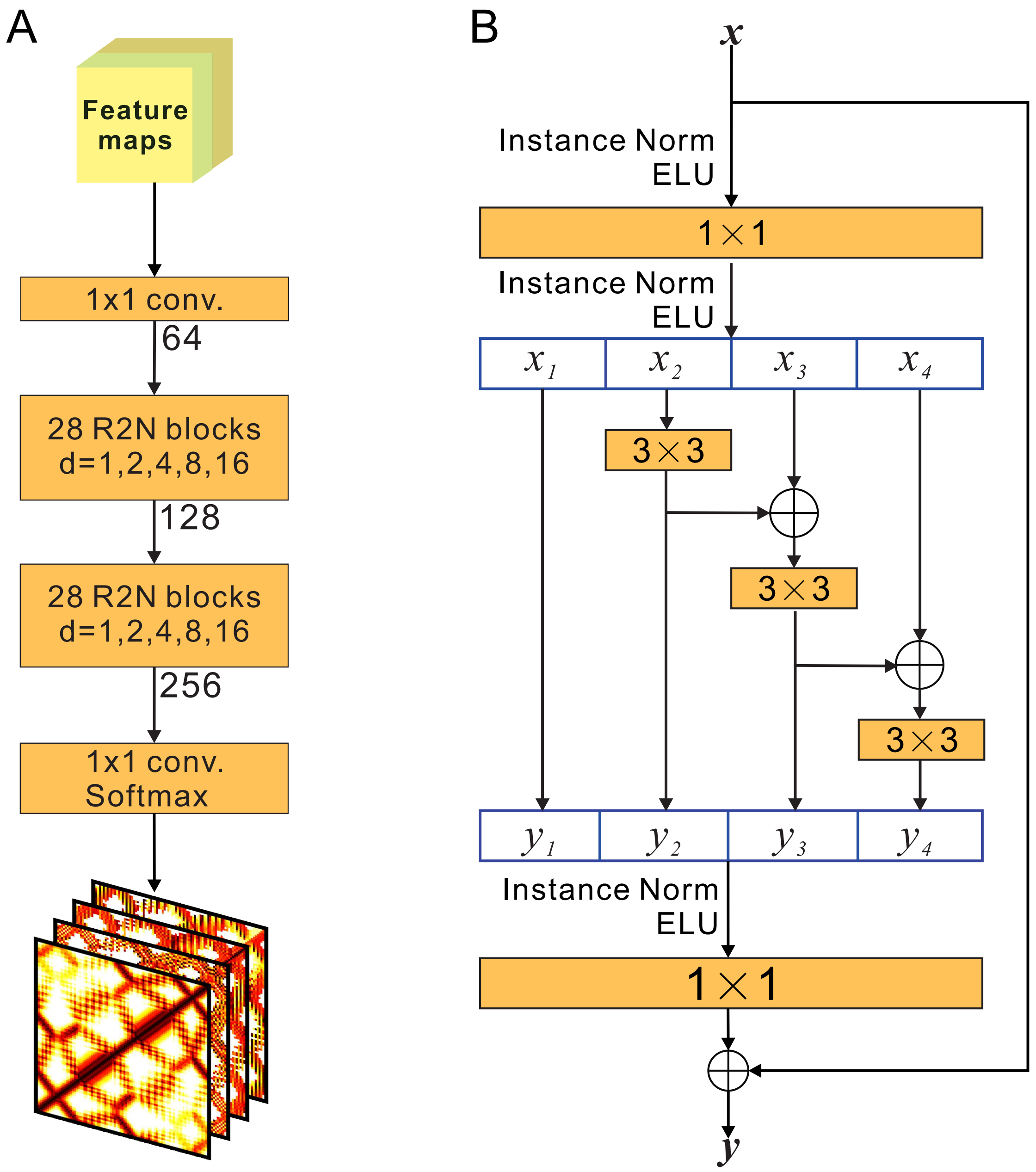


**Figure S3. Architecture of neural network used in trRosettaX-Single.** A. the architecture of Res2Net_Single. We employ dilated convolutions with different dilation rates (denoted by d). B. a basic Res2net (R2N) block in Res2Net_Single.

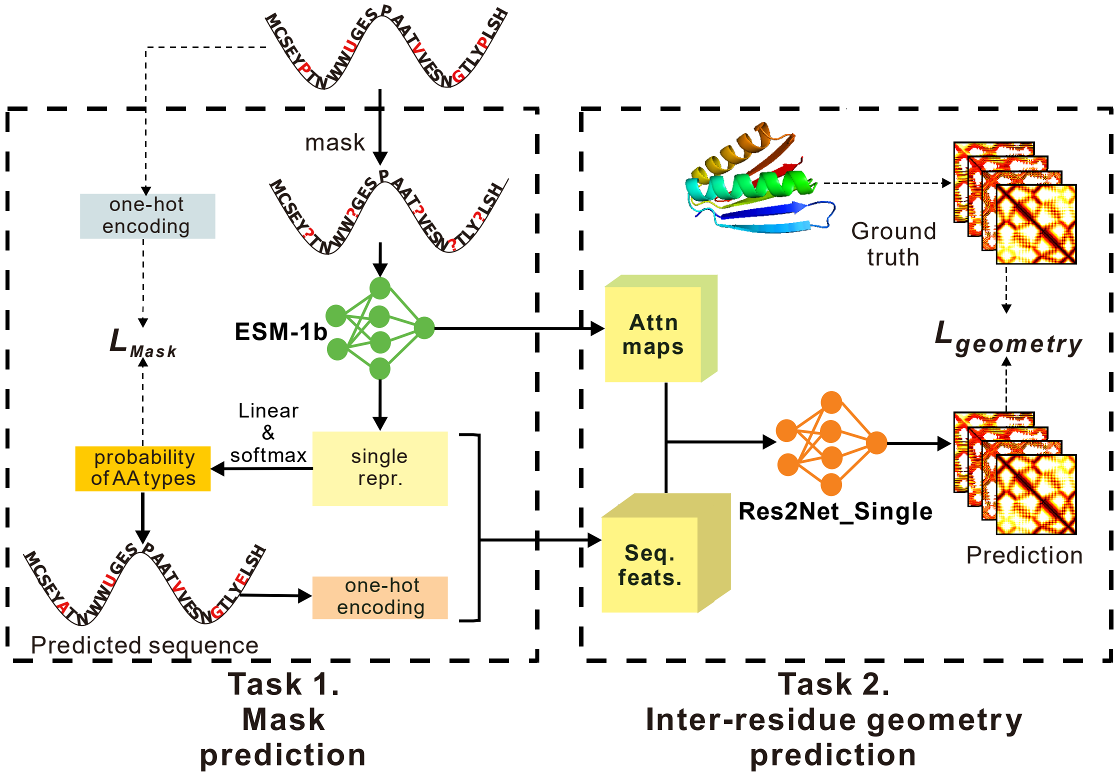


**Figure S4. Development of s-ESM-1b based on supervised re-training of ESM-1b.** We re-train ESM-1b under the supervision of two tasks, starting from its pre-trained parameters. The first is to predict the amino acid types of the randomly masked positions (highlighted in red), supervised by the cross-entropy loss ($L_{mask}$) between the predicted probability distributions and the one-hot encoding of real types. The second is to predict the inter-residue geometry by feeding the sequence representation and attention maps of the masked sequence as well as the one-hot encoding of the predicted sequence into Res2Net_Single, supervised by its cross-entropy loss with the native geometry ($L_{geometry}$). The parameters in Res2Net_Single are also updated here.
